# Spatial moment description of birth-death-movement processes incorporating the effects of crowding and obstacles

**DOI:** 10.1101/267708

**Authors:** Anudeep Surendran, Michael J. Plank, Matthew J. Simpson

**Affiliations:** School of Mathematical Sciences, Queensland University of Technology, Brisbane Queensland, Australia; School of Mathematics and Statistics, University of Canterbury, Christchurch, New Zealand; Te Punaha Matatini, A New Zealand Centre of Research Excellence, Auckland, New Zealand.

**Keywords:** Collective cell migration, Spatial moment dynamics, Individual based model, Cell proliferation, Cell migration

## Abstract

Birth-death-movement processes, modulated by interactions between individuals, are fundamental to many cell biology processes. A key feature of the movement of cells within *in vivo* environments are the interactions between motile cells and stationary obstacles. Here we propose a multi-species model of individual-level motility, proliferation and death. This model is a spatial birth-death-movement stochastic process, a class of individual-based model (IBM) that is amenable to mathematical analysis. We present the IBM in a general multi-species framework, and then focus on the case of a population of motile, proliferative agents in an environment populated by stationary, non-proliferative obstacles. To analyse the IBM, we derive a system of spatial moment equations governing the evolution of the density of agents and the density of pairs of agents. This approach avoids making the usual mean-field assumption so that our models can be used to study the formation of spatial structure, such as clustering and aggregation, and to understand how spatial structure influences population-level outcomes. Overall the spatial moment model provides a reasonably accurate prediction of the system dynamics, including important effects such as how varying the properties of the obstacles leads to different spatial patterns in the population of agents.

## 1 Introduction

Movement, birth and death processes are important individual-level mechanisms that can influence population-level outcomes in both biological and ecological systems. Cell migration and cell proliferation, modulated by interactions among neighbouring cells and obstacles, are essential for development (Kurosaka and Kashina 2008), repair (Martin 1997) and disease (Friedl and Wolf 2003). Similarly, in many ecological systems, the movement of individuals, and interactions between individuals, can have important population-level consequences. The emergence of spatial structure in predator-prey systems and plant communities are direct results of interactions between individuals (Tobin and Bjornstad 2003; Law and Dieckmann 2000). These common observations from different areas of the life sciences suggest a role for individual-level, mathematical models to represent molecules, cells, plants and animals. Popular modelling frameworks include lattice-based models and continuous space lattice free models (Plank and Simpson 2012; Dyson and Baker 2015). Some of these models consider single species populations (Lewis 2000; Middleton et al. 2014), while others incorporate the influence of interactions among different subpopulations (Jin et al. 2018; Murrell 2005; Smith et al. 2017).

Cell migration in living tissues involves complicated heterogeneous environments that are occupied by various biological structures and scaffolds of varying size, shape and adhesive properties (Ellery et al. 2014; Ellery et al. 2016). Such obstacles can have a significant impact on the migration of cells due to the interplay between crowding and cell-to-substrate adhesion (Welch 2015; Sun and Zaman 2017; Simpson and Plank 2017). The extracellular matrix (ECM), composed of polysaccharides and proteins, is an example of a biological obstacle which influences the cell migration in many different ways, such as providing biochemical stimuli, mechanical cues and steric hindrances (Zaman et al. 2006; Harley et al. 2008). ECM geometry can regulate the motility of cancer (Condeelis and Segall 2003) and immune cells (Bajenoff et al. 2006). The highly compartmentalised structure of the cytoplasm and the presence of macromolecular obstacles such as nucleic acids and proteins within intracellular environments can have a significant impact on biochemical reactions (Tan et al. 2013; Hansen et al. 2016) and physical transport processes inside the cell (Ghosh et al. 2016; Smith et al. 2017). These examples from various biological organisational levels suggest that the incorporation of both obstacles and their crowding effects in a mathematical model of cell migration is important.

Standard models of biological and ecological systems are based on the mean field assumption which, roughly speaking, assumes that individuals in the population encounter each other in proportion to their average density (Law and Dieckmann, 2000). Classical examples of mean field models include Lotka-Volterra models of ecological competition (Murray 1989), the Keller-Segel model of chemotaxis (Keller and Segel 1971), the logistic growth model to describe population dynamics (Edelstein-Keshet 2005) and the Fisher-Kolmorogov model to describe wound healing (Johnston et al. 2015). Standard mathematical models based on ordinary and partial differential equations routinely invoke the mean field assumption. In this work we take a more general approach by accounting for spatial correlations by developing continuous descriptions of the dynamics of individuals, dynamics of pairs of individuals, and so on (Plank and Law, 2015). In general this approach leads to an infinite hierarchy of equations governing the spatial moments of the system which we approximate using a moment closure assumption (Murrell et al. 2004; Raghib et al. 2011).

The first studies that used spatial moments focused on modelling birth-death processes in ecology with a single species (Bolker and Pacala 1997; Lewis 2000), and later studies examined competition and prey-predator interactions in a multi-species birth-death framework (Murrell 2005; Barraquand and Murrell 2013). More recently these ecological models have been extended to include movement processes that are motivated by observations from cell biology experiments (Baker and Simpson 2010; Simpson et al. 2013). However, these first models that include movement are lattice-based, which means that the movement of individuals is restricted to an artificial lattice. Lattice-free moment dynamics models of cell migration and cell proliferation are much more recent (Middleton et al. 2014; Binny et al. 2015, 2016a, 2016b). Unlike the lattice-based models in which volume exclusion is strictly enforced, lattice-free models use interaction kernels to give a more realistic description of interactions because cells are able to deform as they move close to neighbouring cells (Le Clainche and Carlier 2008). In this work we present a lattice-free model of cell migration, cell proliferation and cell death. In an attempt to make the model relevant to *in vivo* conditions, we take a multi-species approach so that we can consider one species to be a population of motile and proliferative cells, whereas the other population represents an immobile subpopulation of obstacles.

## 2 Individual-based model

A key feature of the individual-based model (IBM) is that we consider the total population to be composed of several different subpopulations. This gives great flexibility since the model can be used to study the movement, proliferation and death of different types of individuals, and it also allows for different types of interactions between the different subpopulations. The state of the IBM depends on the position of each individual, and the properties of agents within each subpopulation. In the model, we have a total of *N(t)* agents that are associated with *I* subpopulations. The location of the *n^th^* agent is *x_n_* ∈ ℝ^2^, and each agent belongs to a particular subpopulation, *i_n_* ∈ {1, 2,…, *I*}, where *n* = 1, 2,…, *N(t)*. We always initiate the IBM with individuals distributed according to a spatial Poisson process, meaning that the IBM is relevant to spatially homogeneous problems without macroscopic gradients in density of individuals (Jin et al. 2018).

Individuals undergo movement, proliferation and death events with a rate per unit time given by 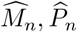 and 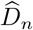 respectively. The IBM is a continuous-time Markov process, where the probability of agent n undergoing a movement during a short time interval, of duration *δt*, is 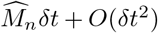. Similar expressions govern the probability of proliferation and death events occurring within a short time interval. The total event rates are comprised of a neighbour independent, intrinsic component, and a neighbour-dependent component accounting for interactions with neighbouring individuals. The neighbourhood contribution is specified by an interaction kernel that depends upon the distance, |*ξ*|, between pairs of individuals. We assume the interaction kernel is isotropic and decays to zero at larger distances, |*ξ*|, ensuring that only relatively close individuals interact. For movement events, the intrinsic component of the movement rate of an individual from subpopulation *i* is denoted by *m_i_*, and the interactions from a neighbouring agent from subpopulation *j* is governed by an interaction kernel 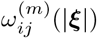. Hence we write the net movement rate of individual *n* as,

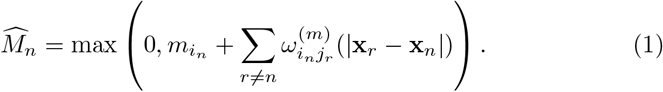

Here, *i_n_* = 1 if individual *n* is from the first subpopulation, and *i_n_* = 2 if individual *n* is from the second subpopulation, and so on. Similarly, we write the net proliferation and net death rates as,

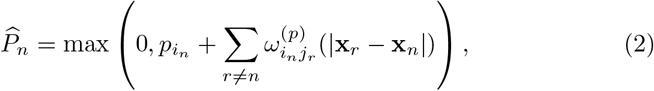

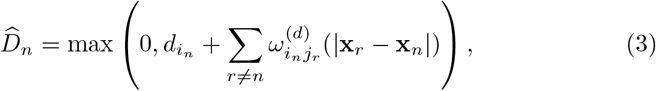

where *p_i_* and *d_i_* are the intrinsic proliferation and death rates, respectively, for an individual from subpopulation *i*. For simplicity, we assume a constant death rate and a fixed distribution for the direction of placement of daughter agents which is unaffected by the interactions with other individuals.

When an individual from subpopulation *i* undergoes a movement event, it travels a displacement *ξ* that is drawn from a probability density function (PDF) 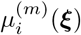. If an individual from subpopulation *i* proliferates, a daughter of the same subpopulation is placed at a displacement *ξ*, that is drawn from a PDF 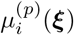. We refer to the placement of daughter agents as a result of a successful proliferation event as agent *dispersal.* Now we generalise the movement displacement PDF by introducing a neighbour-dependent bias vector, to accommodate the influence of neighbouring individuals upon the direction of movement. We introduce an interaction kernel, 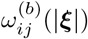, to account for neighbour-dependent directional bias acting on the reference individual, from subpopulation *i*, due to the presence of a second individual, from subpopulation *j*, at a displacement ξ. The neighbour-dependent bias is defined as the gradient of interaction kernel, 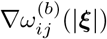. This definition is same as that of Binny et al. (2016b), however here we generalise that previous model to account for the contributions to the directional bias arising from multiple subpopulations. The net bias vector is the sum of contributions from each of the neighbouring individuals, and a constant neighbour-independent global bias 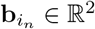, giving

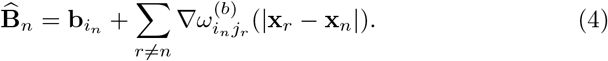

The angular direction of the net bias vector, denoted by 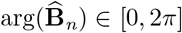, is the preferred direction of movement for a particular individual. In this framework, the preferred direction of movement is driven, in part, by the sum of the gradient of the interaction kernels. The strength of bias is given by the magnitude of bias vector, 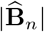. Now we assume that the neighbouring individuals affect the direction of movement arg(*ξ*) ∈ [0, 2*π*], but not the actual distance moved, |*ξ*|.

During a movement event, the direction of movement is drawn from a von Mises distribution, 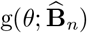 whose concentration parameter is 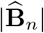, and the mean direction is given by 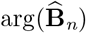,

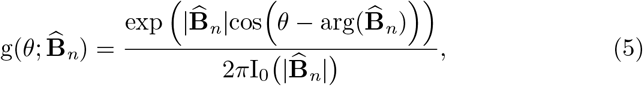

where *I*_0_ is the zero order modified Bessel function. This PDF ensures that individuals are most likely move in the direction of 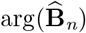, and the bias to move in this preferred direction increases with 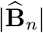. When the net bias is zero, 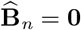, the von Mises distribution reduces to the uniform distribution (Binny et al. 2016b). Individuals located where the gradient is steep will have a large 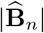, and are more likely to move in the direction of 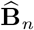. Individuals located where the gradient is relatively flat will have a weaker bias and the direction of movement becomes almost uniformly distributed (Browning et al. 2017). We assume that the distance moved by an agent is independent of local crowding, and is given by a fixed PDF, *u_i_*(|*ξ*|). Hence the net movement displacement PDF is the product of the distance PDF and the direction PDF, giving

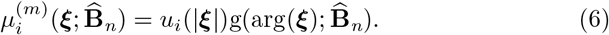

### 2.1 IBM for motile, proliferative agents and stationary obstacles

The IBM can be applied to numerous problems involving populations composed of various combinations of motile and stationary subpopulations by appropriate choice of parameters. Here we focus on an important scenario with two distinct subpopulations, *I* = 2. We consider the first subpopulation, *i* = 1, to be a group of agents undergoing movement, proliferation and death events. This first subpopulation can be thought of as a population of motile, proliferative cells. The first subpopulation interacts with a second subpopulation, *i* = 2, that is composed of stationary, non-proliferating obstacles. The event rates for individual agents in the first subpopulation will be influenced by the presence of both obstacles and other agents in their neighbourhood, given by Equations (1)–(3). Since the obstacles never undergo birth, death or movement events, they contribute to the overall dynamics by interactions between the obstacles and the agents.

We introduce interaction kernels: 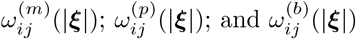, to account for the contribution from surrounding agents and obstacles to the movement rate, proliferation rate, and the directional bias of an agent, respectively. We choose these interaction kernels to be two-dimensional Gaussian functions. The movement interaction kernel is given by,

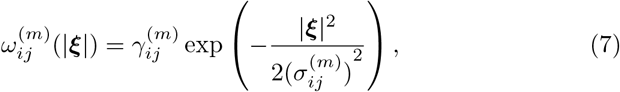

where 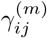 and 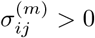 represent the interaction strength and the spatial extent of interaction, respectively. We assume a similar form for the proliferation and bias kernels, given by,

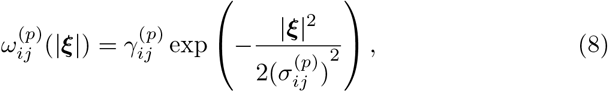

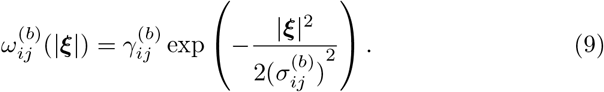

In our simulations we have two subpopulations. The first subpopulation, denoted *i* = 1, corresponds to the motile and proliferative agents. The second subpopulation, denoted *i* = 2, corresponds to the stationary, non-proliferative obstacles. The intrinsic rates of movement and proliferation of agents are *m*_1_ and *p*_1_, respectively. These rates for obstacles are zero. The presence of subpopulations of agents and obstacles results in different types of interactions. The interactions involving the pairs of individuals of same type such as, agent-agent and obstacle-obstacle pairs, are specified by 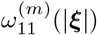 and 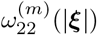, respectively. Similar kernels apply for proliferation and bias. Interactions involving individuals from different subpopulations are specified by the interaction kernels 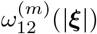 and 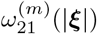. Since obstacles are both stationary and non-proliferative, the presence of neighbouring agents and obstacle does not affect the dynamics and spatial arrangement of obstacles in any way. Hence the interaction strengths 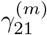 and 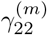 are both set to zero. Positive 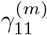 and 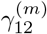 values enhance the movement rate of agents, and can be thought of as representing contact stimulation of migration. In contrast, a negative interaction strength results in a reduction of the movement rate, which can be thought of as representing contact inhibition of migration. Similarly, the net proliferation rate of agents, and the nature of directional bias depend on the sign of interaction strength.

We use a univariate Gaussian distribution, with mean 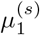 and standard deviation 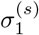, to specify the distribution of movement distance for agents *u_1_*(|*ξ*|). The PDFs of movement displacement, 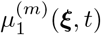 and 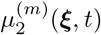, also require the specification of the movement distance distribution, *u_1_*(|*ξ*|), and the neighbour-dependent direction probability density function given by Equation (5). For simplicity, the PDF for the dispersal of daughter agents arising from proliferation events, 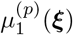, is chosen to be neighbour independent, and specified as a bivariate Gaussian distribution with zero mean and standard deviation 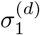.

### 2.2 Numerical implementation

We simulate the IBM using the Gillespie algorithm (Gillespie 1977) implemented in FORTRAN. In each simulation the population is initially composed of *N*_1_ (0) agents and *N*_2_ (0) obstacles, distributed according to a spatial Poisson process across a square domain of size *L × L*. The movement, proliferation and death rates of the agents are computed using Equations (1)–(3). For this study, we use a constant death rate for all the agents, hence the neighbour-dependent term in Equation (3) is set to zero. The sum of event rates of all agents is given by,

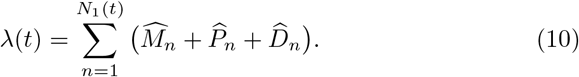

Since the event rates for obstacles are always zero, those terms do not contribute to λ(*t*). The time interval between consecutive events is exponentially distributed with mean 1/λ(*t*). At each event time, one of the three possible events occurs to an agent. The probability of occurrence of an event is proportional to the rate of that event. For a movement event, the agent moves a displacement specified by the bias vector and movement displacement PDF in Equation (4) and Equation (6), respectively. For a proliferation event, the proliferative agent disperses a daughter agent at a displacement specified by the PDF 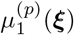, and the total number of agents increases by one. For a death event the total number of agents is reduced by one.

To provide a mathematical description of the IBM we analyse the dynamics of the first and second spatial moments of the agents and obstacles. The first moment of agents and obstacles is given by dividing the total number of agents and obstacles by the area of the domain, giving *N*_1_(*t*)/*L*^2^ and *N*_2_(*t*)/*L*^2^, respectively. We use a pair correlation function (PCF) to quantify the second spatial moment. Here the PCF depends on both the separation distance, *r*, and time, *t* (Angew et al. 2014; Dini et al. 2018). However, to be consistent with previous studies, our notation focuses on the separation distance, *r*, only. Since there are two different subpopulations, we will have two different PCFs. First, we denote the auto-correlation PCF between agents as *C*_11_(*r*). Second, we denote the cross-correlation between agents and obstacles as *C*_12_(*r*). To compute the auto-correlation function, we consider a reference agent at x_*i*_, and calculate all distances, *r* = |x_*j*_ – x_*i*_|, to the other *N*_1_ – 1 agents. We follow the same procedure with each of the remaining agents until all agents have acted as the reference agent. Note that we always take care to measure distances across periodic boundaries. With this information, the auto-correlation PCF is constructed by enumerating the distances between pairs of agents that fall into the interval, [*r* – *δr*/2, *r* + *δr*/2]. That means we use a bin width of *δr*. To ensure that *C*_11_(*r*) = 1 in the absence of spatial structure, we normalise the bin count by a factor of *N*_1_(*t*)(*N*_1_(*t*) – 1)(2*πrδr*)/*L*^2^. When *C*_11_(*r*) > 1, we have a larger number of pairs of agents separated by a distance r than we would have in the spatially random population. In contrast, for *C*_11_(*r*) < 1, we have a smaller number of pairs of agents separated by a distance r than we would have in the spatially random population. Similarly, we compute the crosscorrelation PCF, *C*_12_(*r*), by counting, binning and normalising all distances between agents and obstacles. A similar calculation could be made for *C*_22_(*r*), by counting, binning and normalising all distances between pairs of obstacles. However, since obstacles are stationary, non-proliferative and initialised at random, we always have *C*_22_(*r*) = 1. Only when *C*_11_(*r*) = *C*_12_(*r*) = *C*_22_ = 1 are agents and obstacles are arranged at random, which is an implicit assumption in all mean-field models. One of the important features of our model and our analysis is that we can have spatial structure, such as clustering or regular spatial patterns, present in the population. This is signified by having *C*_11_(*r*) ≠ 1 and *C*_12_(*r*) ≠ 1.

There are several key variables relevant to specifying different obstacle fields and different obstacle properties. These include obstacle density, obstacle size and obstacle interactions, such as whether obstacles are adhesive or repulsive. We will explore how systematically varying these properties influences the development of spatial structure in a population of motile and proliferative agents that are placed into an environment containing obstacles. We first explore this by performing repeated realizations of the IBM and analysing averaged ensemble data in terms of the first spatial moment and PCF. Second, we then compare these results with the prediction from our analysis of spatial moment dynamics in Section 5. A summary of key variables and notation for these calculations is given in Table 1.

**Table 1.** Model parameters and typical values.

| Parameter | Symbol | Value |
| --- | --- | --- |
| **Densities** |  |  |
| Agents | $Z_{1,1}(t)$ | 0.25 to 0.9 |
| Obstacles | $Z_{1,2}(t)$ | 0.0 to 0.375 |
| Initial density of agents | $Z_{1,1}(0)$ | 0.25 |
| Initial density of obstacles | $Z_{1,2}(0)$ | 0.0 to 0.375 |
| **Intrinsic rates** |  |  |
| Birth | $p_1, p_2$ | 1, 0 |
| Death | $d_1, d_2$ | 0.5, 0 |
| Movement | $m_1, m_2$ | 5, 0 |
| **Neighbour-dependent interaction strengths** |  |  |
| Birth | $\gamma_{11}^{(p)}, \gamma_{12}^{(p)}, \gamma_{21}^{(p)}, \gamma_{22}^{(p)}$ | -0.38 to 0 |
| Movement | $\gamma_{11}^{(m)}, \gamma_{12}^{(m)}, \gamma_{21}^{(m)}, \gamma_{22}^{(m)}$ | 0 |
| Bias | $\gamma_{11}^{(b)}, \gamma_{12}^{(b)}, \gamma_{21}^{(b)}, \gamma_{22}^{(b)}$ | -0.2 to 0.4 |
| **Spatial extent of interactions** |  |  |
| Birth | $\sigma_{11}^{(p)}, \sigma_{12}^{(p)}, \sigma_{21}^{(p)}, \sigma_{22}^{(p)}$ | 0.25 to 0.6 |
| Movement | $\sigma_{11}^{(m)}, \sigma_{12}^{(m)}, \sigma_{21}^{(m)}, \sigma_{22}^{(m)}$ | 0.25 to 0.6 |
| Bias | $\sigma_{11}^{(b)}, \sigma_{12}^{(b)}, \sigma_{21}^{(b)}, \sigma_{22}^{(b)}$ | 0.25 to 0.6 |
| **Movement and dispersal distance** |  |  |
| Mean movement distance | $\mu_1^{(s)}, \mu_2^{(s)}$ | 0.4 |
| Standard deviation movement distance | $\sigma_1^{(s)}, \sigma_2^{(s)}$ | 0.1 |
| Standard deviation dispersal distance | $\sigma_1^{(d)}, \sigma_2^{(d)}$ | 0.5 |

## 3 Continuous description

Here, we derive a continuum approximation for the IBM in terms of spatial moments. To keep our work as general as possible, we present the derivation for an arbitrary number of motile and proliferative subpopulations. Then we present a specific case of the continuous description where there are two subpopulations: the first subpopulation is a population of motile and proliferative agents and the second subpopulation is a population of stationary and non-proliferative obstacles. As previously described, we consider a spatially homogeneous environment. This means that the probability of finding an individual in a given small region is independent of the position of that small region. Hence the key quantity of interest is the displacement between individuals. The first spatial moment, *Z*_1,*i*_(*t*), is the average density of individuals from subpopulation *i*. The second spatial moment, *Z*_2,*i,j*_(*ξ,t*), is the average density of pairs of individuals, consisting of an individual from subpopulation *j* at a displacement *ξ* from an individual belonging to subpopulation *i*. The third spatial moment, *Z_3,i,j,k_*(*ξ, ξ′,t*), is the average density of triplets of individuals. Here, this triplet consists of an individual from subpopulation *j* at displacement from an individual belonging to subpopulation *i*, and an individual from subpopulation *k* at displacement *ξ′* from the individual belonging to subpopulation *i*. Mathematical definitions of these spatial moments are presented in Section 1 of the Supplementary Material.

### 3.1 Dynamics of spatial moments

The expected rates of movement and proliferation of an individual, denoted *M*_1*i*_(*t*) and *P*_1*i*_(*t*), depend on the contribution from another individual at a displacement ξ. The conditional probability of having an individual from subpopulation *j* located at *x* + *ξ* given that an individual from subpopulation *i* is located at *x*, is *Z_2,ij_*(*ξ,t*)/*Z*_1*i*_(*t*). The derivation of the conditional probability in terms of the spatial moments is provided in Section 2 of the Supplementary Material. The expected movement and proliferation rates of an individual from subpopulation *i* is given by multiplying this conditional probability by the corresponding interaction kernels, 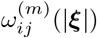 or 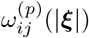, respectively, and integrating over all possible displacements. These details are outlined in Section 3 of the Supplementary Material, and lead to

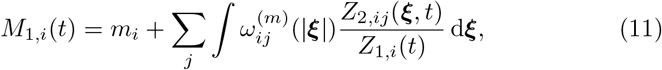

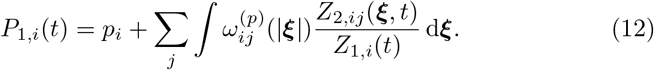

We now consider the dynamics of the first moment. The first moment dynamics depends solely on the balance between the expected rate of proliferation, *P*_1,*i*_(*t*), and death rate, *d_i_*. The movement and neighbour-dependent directional bias does not directly influence the dynamics of the first moment. The time evolution of the first moment is given by,

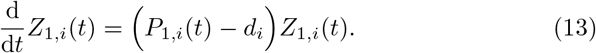

The dynamics of the average density of pairs of agents depends on the conditional occupancy of a third agent in the neighbourhood. The conditional probability of having an individual from subpopulation *k* located at *x* + *ξ*′ given that an individual from subpopulation *i* is located at *x* and an individual from subpopulation *j* located at *x* + *ξ*, is *Z_3,i,j,k_*(*ξ, ξ′,t*)/*Z_2,i,j_*(*ξ,t*). The expected movement and proliferation rates for an individual from subpopulation *i*, which forms a pair with an individual from subpopulation *j*, separated by a displacement *ξ*, denoted by *M_2,i,j_*(*ξ,t*) and *P_2,i,j_*(*ξ,t*), respectively. These rates depend on the conditional presence of another individual from subpopulation *k* at a displacement *ξ′*. Hence, the expected rates are found by multiplying *Z_3,i,j,k_*(*ξ,′,t*)/*Z_2i,j_*(*ξ,t*) by the interaction kernels and integrating over all possible displacements as follows,

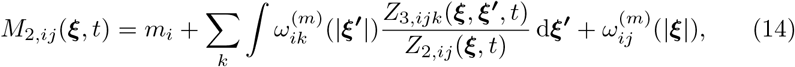

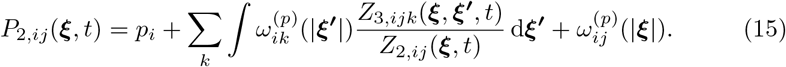

The third term on the right of Equations (14)–(15) accounts for the direct influence of the individual from subpopulation *j* at a displacement *ξ* from the individual from subpopulation *i*.

The gradient of the interaction kernel specifies the contribution of other individuals to the bias vector. The expected net bias vector for an individual from subpopulation *i*, conditional on presence of an individual from subpopu-lation *j*, is given by,

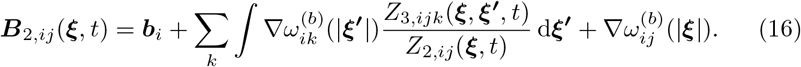

Again, the third term on the right of Equation (16) accounts for the direct influence of an individual from subpopulation *j*, at a displacement *ξ*, from the individual belonging to subpopulation *i*. We note that Equation (16) combines directional bias (Binny et al. 2016a, 2016b) with multi-species spatial moment equations (Law and Dieckmann 2000; Murrell 2005; Plank and Law 2015) in a way that has not been considered previously. Now we consider the PDF for the movement displacement *ξ′* for an individual from subpopulation *i* conditional on the presence of an individual from subpopulation *j* at displacement *ξ* as,

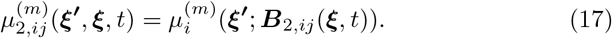

For the dynamics of the second moment, we must consider two factors: (i) the loss of pairs of agents at displacement *ξ*; and (ii) the creation of pairs at displacement *ξ*. The loss of pairs occurs either by movement or death events, whereas the creation of pairs occurs through movement or proliferation events. The time evolution of the second moment is given by,

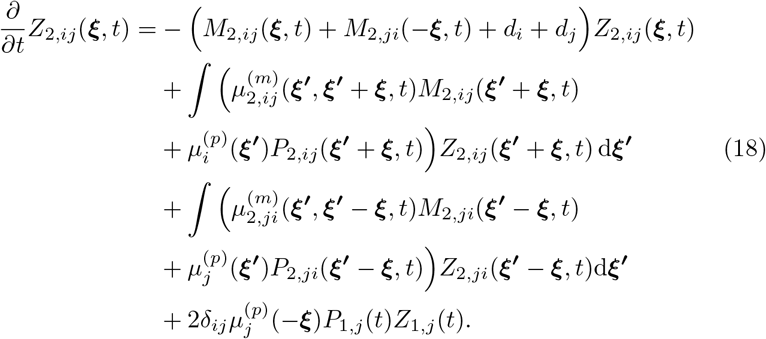

In Equation (18), the two integral terms and the factor of two in the last term on the right arises due to the fact that a pair can be created or destroyed by either of the individuals in that pair. A detailed description of the derivation of the dynamics of the second spatial moment is provided in Section 4 of the Supplementary Material.

Just as the dynamics of the first moment depends on the second moment, we see that the dynamics of the second moment depends on the third moment. If we continue in this way we could develop an infinite hierarchy of equations which is difficult to analyse (Ovaskainen and Cornell 2006; Finkelshtein et al. 2009; Ovaskainen et al. 2014). However, previous studies focusing on applications in cell biology (Binny et al. 2016a, 2016b) and ecology (Law and Dieckmann 2000) provide useful results by closing the infinite system of moment equations to produce an approximate truncated system. Here we follow a similar approach and approximate the third order terms in Equations (14)–(16) using a moment closure approximation. While various approximations, such as the power-1 closure, power-2 closure and Kirkwood superposition approximation, are available (Murrell et al. 2004), in this study we focus on the power-2 closure scheme (Law et al. 2003) that is given by,

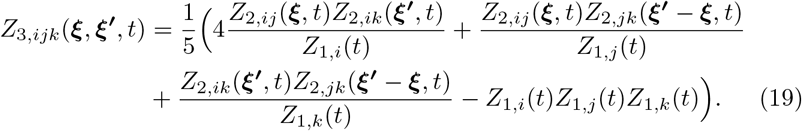

### 3.2 Spatial moment description for motile, proliferative agents and stationary obstacles

For clarity, we now present the governing equations for the specific case of a population of motile, proliferative agents in an environment containing stationary, non-proliferative obstacles. The first spatial moment of agents and obstacles are denoted *Z*_1,1_(*t*) and *Z*_1,2_(*t*), respectively. The second spatial moments, corresponding to the density of pairs are denoted: *Z*_2,11_(*ξ, t*); *Z*_2,12_(*ξ,t*); *Z*_2,21_(*ξ,t*); and *Z*_2,22_(*ξ,t*). The terms *Z*_2,11_(*ξ,t*) and *Z*_2,22_(*ξ,t*) correspond to the average densities agent-agent pairs and obstacle-obstacle pairs. The other two terms, *Z*_2,12_(*ξ,t*) and *Z*_2,21_(*ξ,t*), corresponds to the average densities of agent-obstacle pairs, and obstacle-agent pairs.

The expected rate of movement of an agent, *M*_1,1_(*t*), is given by multiplying the interaction kernels of movement, 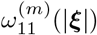 or 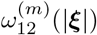, by the conditional probability of having an agent or obstacle present at a displacement *ξ* from the reference agent, and integrating over all possible displacements. These conditional probabilities of either an agent or obstacle located at a displacement are *ξ* given by *Z*_2,11_(*ξ,t*)/*Z*_1,1_(*t*) and *Z*_2,12_(*ξ,t*)/*Z*_1,1_(*t*), respectively. The expected proliferation rate is also calculated in the same way by replacing the movement kernels with proliferation kernels, 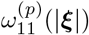 and 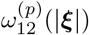. Using this information, the expected rate of movement and proliferation of an agent is given by,

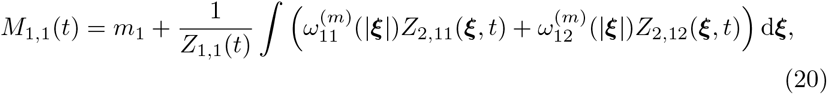

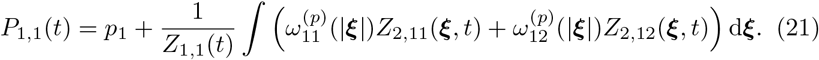

The expected movement and proliferation rates of obstacles *M*_1,2_(*t*) and *P*_1,2_(*t*) are zero.

The time evolution of density of agents depends only on the expected rate of proliferation and death of agents. The density of obstacles remains constant. Hence we have,

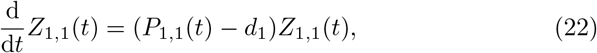

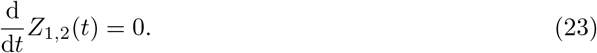

The conditional probability of having an agent located at ξ′, given that an another agent is present at *ξ*, is 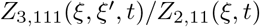. Similarly three more conditional probabilities can be specified by considering different arrangements of agents and obstacles around the reference agent at displacement 0. The expected event rates, *M*_2,11_(*ξ, t*) and *P*_2,11_(*ξ, t*), of an agent conditional on the presence of another agent at displacement *ξ* can be computed by multiplying these conditional probabilities by the corresponding interaction kernels and integrating over all possible displacements. The expected rates are given by,

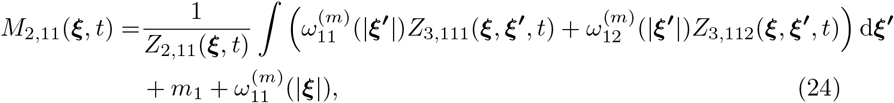

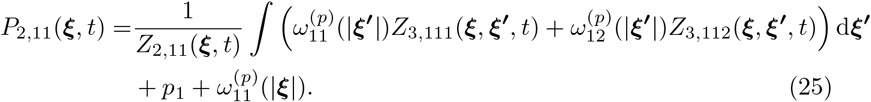

Using similar arguments, we compute the remaining movement and proliferation rates of an agent arising from the presence of an obstacle at a displacement *ξ* as follows,

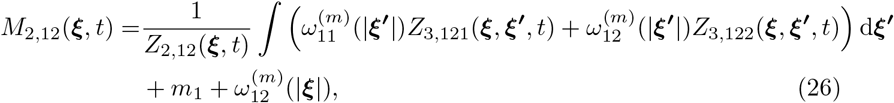

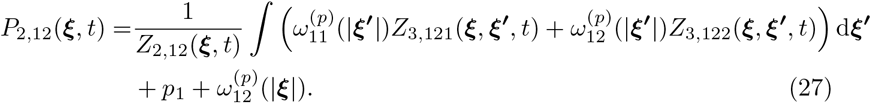

The gradient of the bias kernel gives the contribution of agents and obstacles to the bias vector of the neighbouring agent. The expected net bias vector of an agent conditional on presence of another agent is given by,

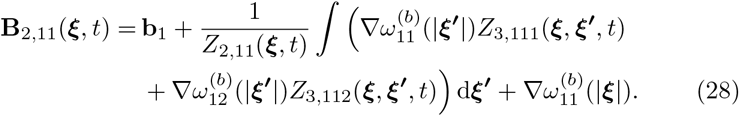

Similarly, the expected net bias vector of an agent conditional on presence of an obstacle is given by,

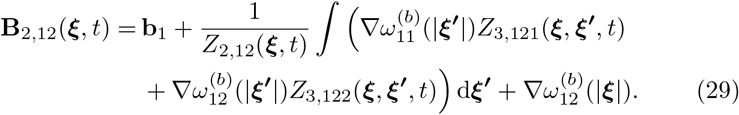

Now we develop the equations governing the dynamics of the second moments. The equations for the density of pairs involving agents depends on the loss of pairs of agents at displacement *ξ*, which can occur either by movement or death, and creation of pair at displacement *ξ* which can occur through movement or proliferation. The density of pairs of obstacles, *Z*_2,22_(*ξ,t*), remains constant over time. The equations governing the dynamics of second moments for the agent-obstacle population are given by,

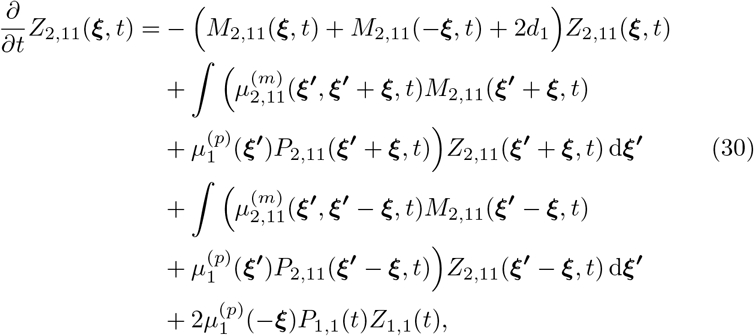

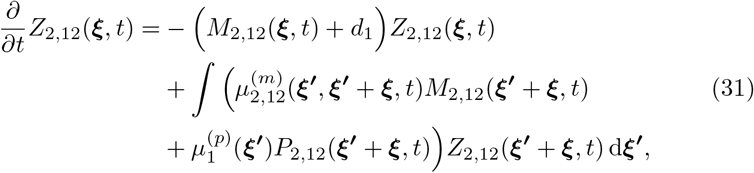

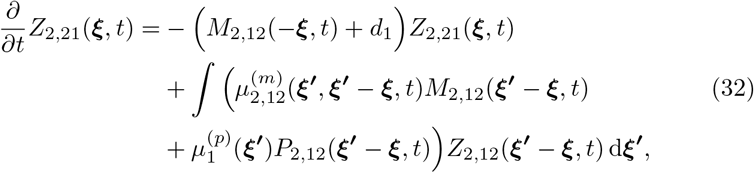

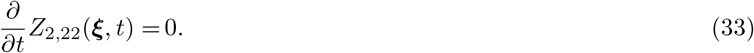

A description of the numerical methods we use to solve Equations (30)–(33) are given in Section 5 of the Supplementary Material.

## 4 Bias landscape

A key feature of the IBM is the neighbour-dependent bias. This formalism helps us understand how the spatial arrangement of obstacles and other agents affects the movement of a particular reference agent. To interpret and visualise the neighbour-dependent bias, we define the *bias landscape* as,

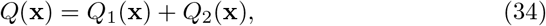

where,

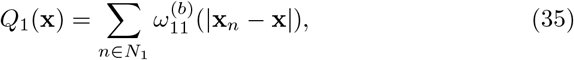

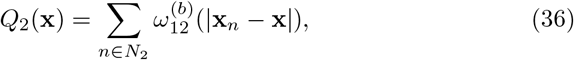

are contributions to overall bias landscape from the agents and the obstacles, respectively. For an agent located at x, *Q*(x) acts as a measure of the degree of crowding. The neighbour-dependent bias vector, **B**_*n*_, is the negative gradient of the bias landscape, –∇*Q*(x). With these definitions it is straightforward to see that agents move in response to the gradient of the bias landscape. Writing the negative gradient of the bias landscape in terms of the contributions from the two subpopulations, –∇*Q*(x) = –∇*Q*_1_(x)–∇*Q*_2_(x), it is clear that the spatial arrangements of both the agents and the obstacles play a role in influencing the movement of agents in the IBM. The incorporation of bias owing to interactions within a particular subpopulation, and interactions from other subpopulations is a very simple, yet powerful feature of our IBM and this can be used to explore a range of behaviours such as enabling the simulation of attraction of individuals within the same subpopulation, and repulsion between pairs of individuals from different subpopulations. This kind of detail, which has never been considered before in the context of a spatial moment model, can be incorporated very simply in our modelling framework.

The influence of bias depends on the spatial arrangement of the agents and obstacles, and properties of the interaction kernels, 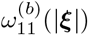 and 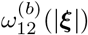, respectively. When the kernels are decreasing functions of |*ξ*|, agents experience a repulsive bias that encourages them to move away from the regions of high crowding. Figure 1 shows how the repulsive bias affects the movement of agents in a particular arrangement. The bias landscape and corresponding level curves due to agent-agent interactions alone are shown in Figure 1(a)-(b). The locations of agents are represented by red dots, and the arrows indicate the preferred direction of movement. The length of these arrows indicates the strength of bias. We note that crowded agents experience a repulsive bias, and prefer to move towards a lower density region. Figure 1(c)-(d) shows the crowding effects generated by obstacles only, and Figure 1(e)-(f) shows how both the agent-agent and agent-obstacle interactions sum to give the total bias landscape. Note that, if the bias interaction strength is negative, then the interaction kernels 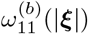 and 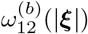 are increasing functions of |*ξ*|. In that case, the orientation of the bias landscape structure would be reversed, and we would have an attractive bias.

**Fig. 1.**
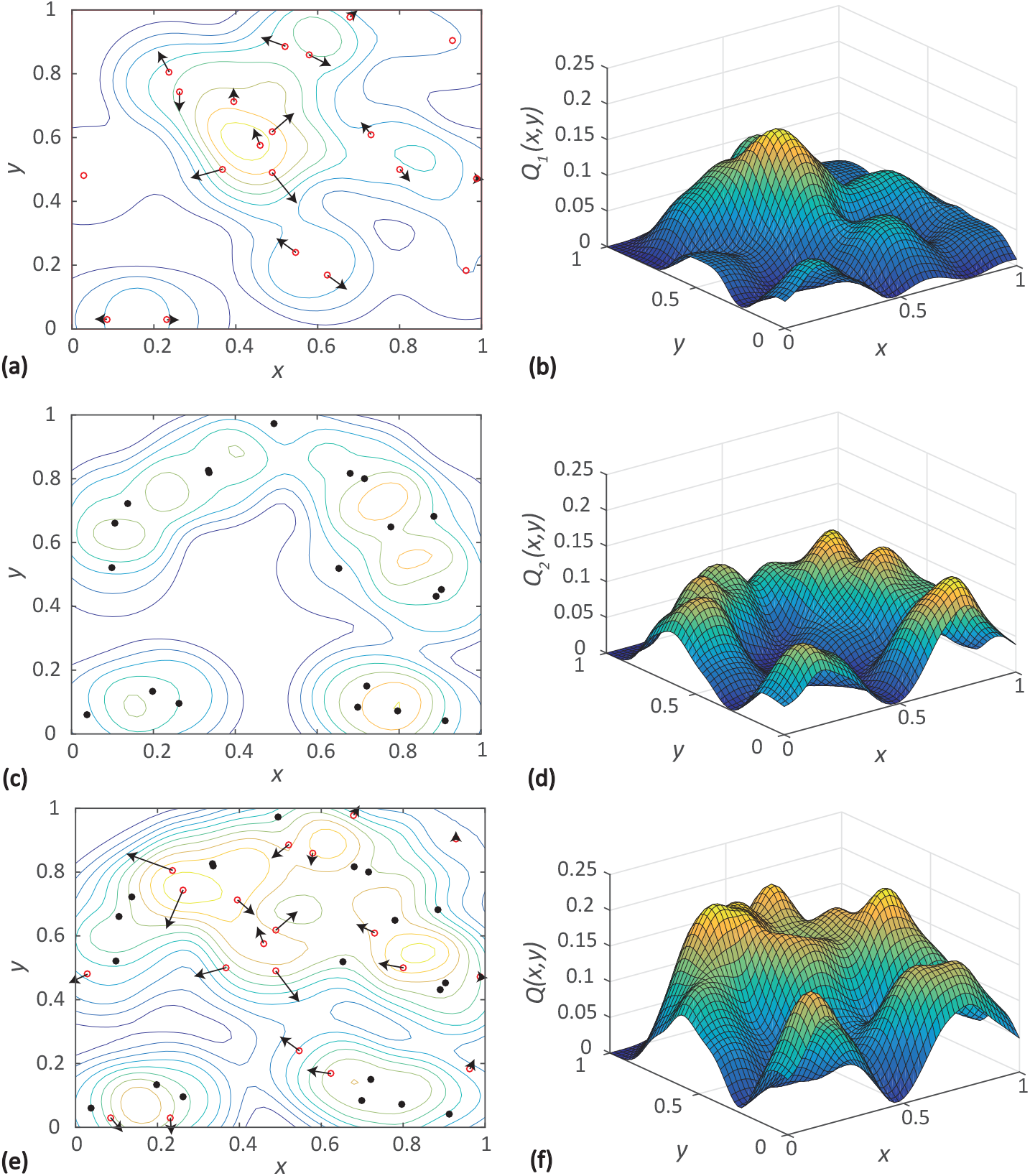
Neighbour-dependent bias visualised for a particular arrangement of 20 agents (red open circles) and 20 obstacles (black dots). In this case we have a positive Gaussian interaction kernel to specify the interactions among agents as well as the interactions between agents and obstacles. **a** Locations of agents (red open circles). **c** Locations of obstacles (black dots). **e** Location of both agents (red open circles) and obstacles (black dots). In each subfigure, **a,c,e**, the locations of individuals are superimposed with the level curves of various components of the bias landscape. **b,d,f** shows the different components of the bias landscape: *Q*_1_(x); *Q*_2_(x); and *Q*(x), respectively. The bias vectors are denoted by black arrows. The bias vectors in a show –∇*Q*_1_(x_*n*_), which is the negative gradient of the component of the bias landscape corresponding to agent-agent interactions. For some agents where the bias vector is sufficiently small we do not show the bias vector. In addition, the bias vectors for stationary obstacles are omitted since the obstacles are stationary. The bias vectors in **e** show –∇*Q*(x_*n*_), which is the negative gradient of the net bias landscape corresponding to the sum of agent-agent and agent-obstacle interactions. The length of arrows indicate the strength of bias. Results in **a-b** correspond to interactions between agents only. Results in **c-d** show interactions between obstacles. Results in **e-f** show the net interactions between agents and obstacles. Parameters are 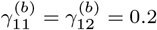 and 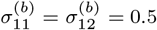.

Another key variable in the IBM is the size of the obstacles, which determines the spatial extent of the obstacle-agent bias. We use the parameter describing the spatial extent of interactions, 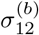, as a proxy for obstacle size. Here we make the natural assumption that larger obstacles correspond to increased 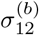, so that larger obstacles tend to exert an influence over a larger neighbourhood. Note that we maintain the bias strength, 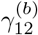, as a constant and we only vary 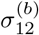 to mimic the influence of obstacle size. Results in Figure 2 show the same spatial arrangement of 20 agents and 20 obstacles as in Figure 1, except that we reduce 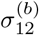 so that the obstacles in Figure 2 influence a smaller neighbourhood than the obstacles in Figure 1. This change in obstacle size does not affect the agent-agent component of the bias landscape, *Q*_1_(x), but it does affect the agent-obstacle component, *Q*_2_(x).

**Fig. 2.**
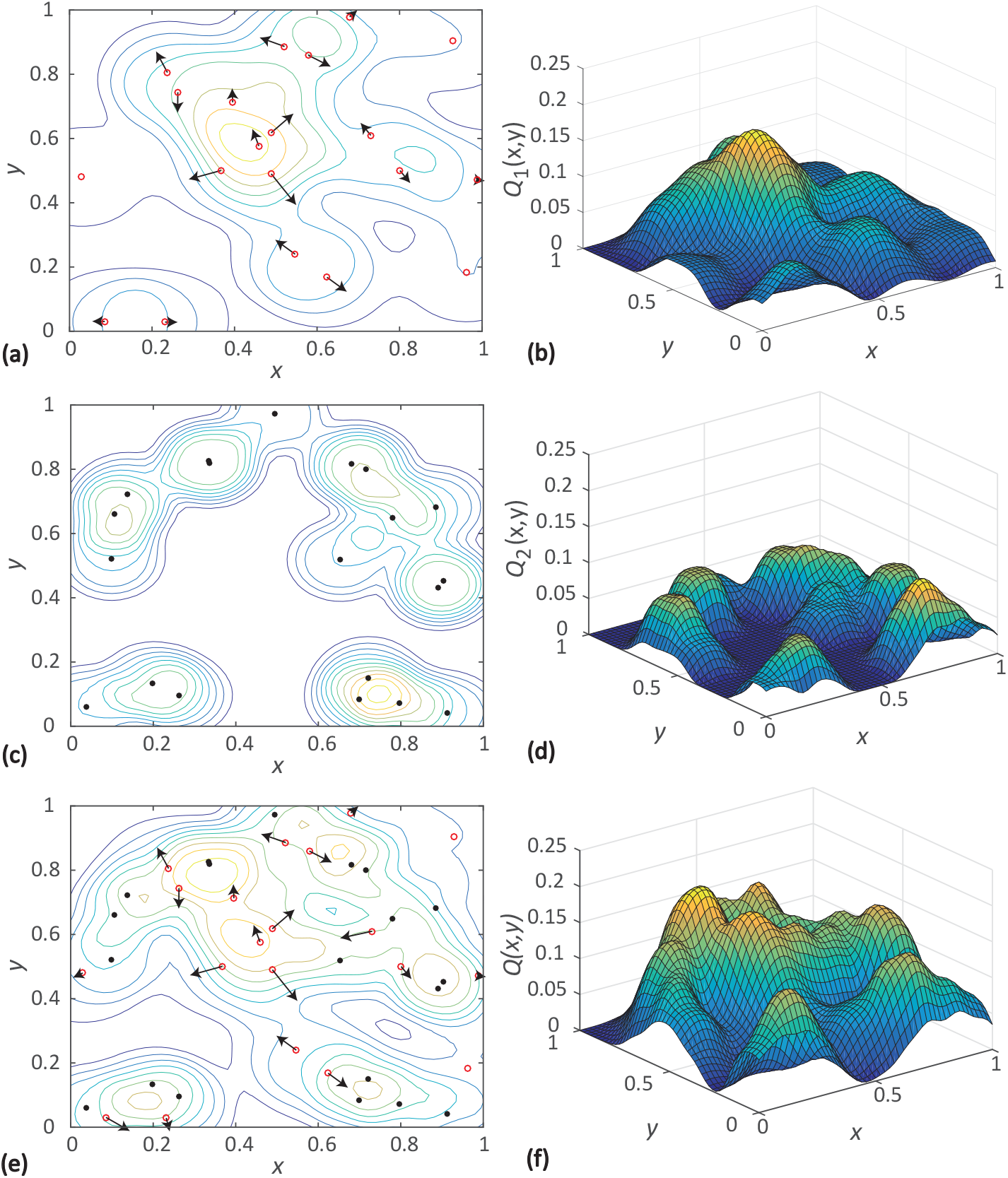
Neighbour-dependent bias visualised for a particular arrangement of 20 agents (red open circles) and 20 smaller obstacles (black dots). In this case we have a positive Gaussian interaction kernel to specify the interactions among agents as well as the interactions between agents and obstacles. **a** Locations of agents (red open circles). **c** Location of obstacles (black dots). **e** Locations of both agents (red open circles) and obstacles (black dots). In each subfigure, **a,c,e**, the locations of individuals are superimposed with the level curves of various components of the bias landscape. **b,d,f** shows the different components of the bias landscape: *Q*_1_(x); *Q*_2_(x); and *Q*(x), respectively. The bias vectors are denoted by black arrows. The bias vectors in a show –∇*Q*_1_(x*_n_*), which is the negative gradient of the component of the bias landscape corresponding to agent-agent interactions. For some agents where the bias vector is sufficiently small we do not show the bias vector. In addition, the bias vectors for stationary obstacles are omitted since the obstacles are stationary. The bias vectors in e show —∇*Q*(x*_n_*), which is the negative gradient of the net bias landscape corresponding to the sum of agent-agent and agent-obstacle interactions. The length of arrows indicate the strength of bias. Results in **a-b** correspond to interactions between agents only. Results in **c-d** show interactions between obstacles. Results in **e-f** show the net interactions between agents and obstacles. Parameters are 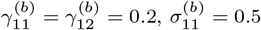 and 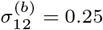.

## 5 Results and discussion

In this section we present snapshots from the IBM to explore the dynamics of the two species agent-obstacle system. In these simulations we systematically vary the density of obstacles, the size of obstacles, and the interactions between agents and obstacles. In addition to presenting IBM simulations, we also present solutions of the spatial moment dynamics model to explore how well the model predicts the dynamics of the different conditions we consider. Another outcome is to examine how various properties of obstacle field influence spatial structure.

### 5.1 Effect of varying the obstacle density

Here we explore how variations in the density of obstacles influence the spatial structure and the dynamics of the agent subpopulation. Results in Figure 3 show a series of snapshots from the IBM. Each row shows snapshots at *t* = 0,10, 20 and 60, whereas each column shows results for different densities of obstacles. The initial number of obstacles is varied from *N*_2_(0) = 0, 50, 100 to 150, in the columns from left-to-right, respectively. In all cases we consider a constant initial number of agents, *N*_1_(0) = 100, and a random initial placement of obstacles and agents. Results in Figure 3(a)-(d) show the most fundamental case where there are no obstacles present, and we note that the multi-species model simplifies to the previous single species model presented by Binny et al. (2016b) in this case. We use this first scenario to emphasise the differences in the population dynamics for the agent subpopulation in presence and absence of obstacles, which are shown in the second, third and fourth column of Figure 3. By repeating the stochastic simulations in Figure 3 many times, we can calculate the density of obstacles, the density of agents, the auto-correlation PCF and the cross-correlation PCF, as shown in Figure 4.

**Fig. 3.**
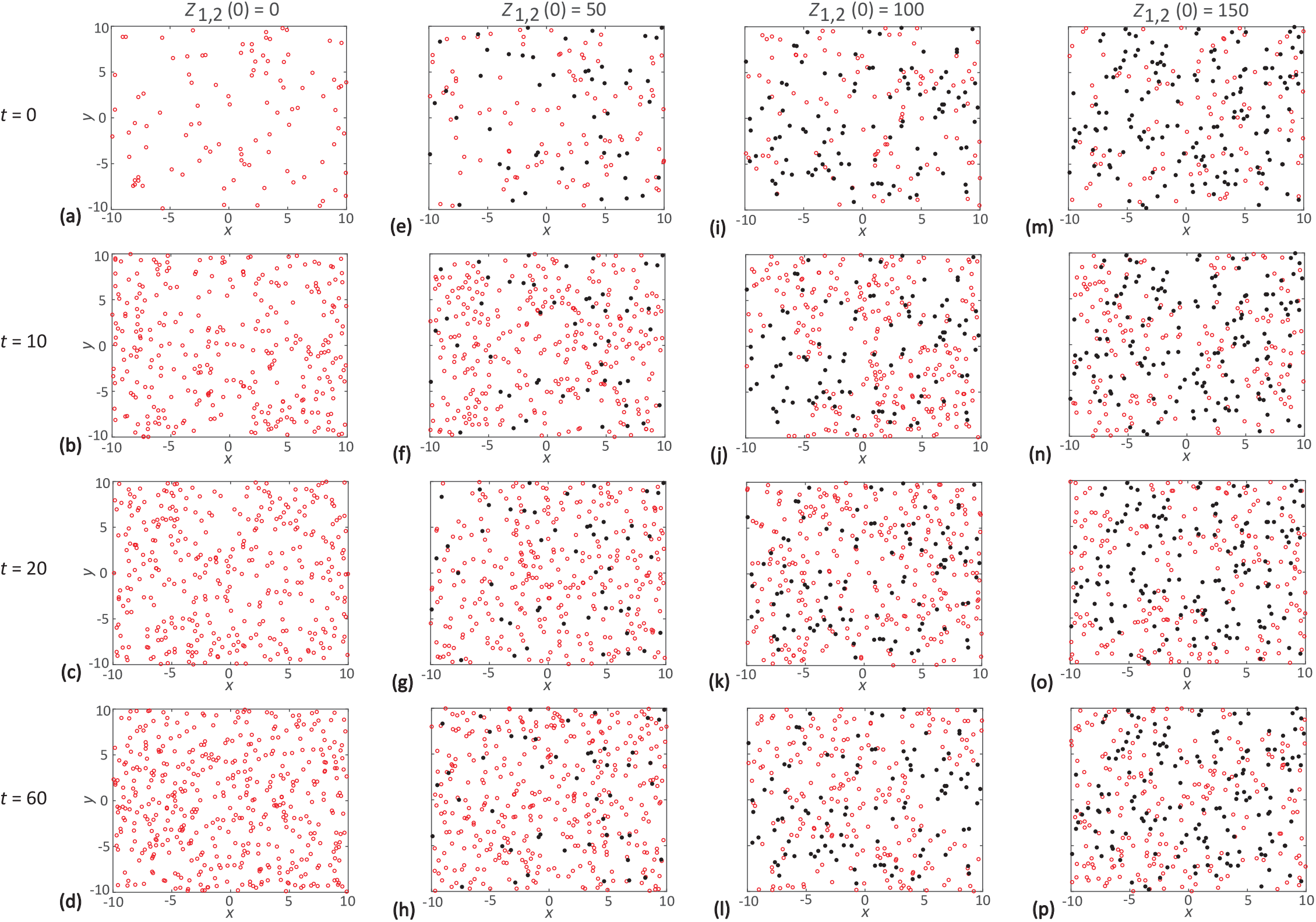
IBM snapshots showing the impact of increasing obstacle density. Each row shows a snapshot of the IBM at *t* = 0, 10, 20 and 60, respectively. Results in **a-d** show the evolution of a population of agents in the absence of obstacles. Results in **e-h, i-l** and **m-p** show the evolution of a population of agents among a population of 50, 100 and 150 obstacles, respectively. In all cases agents are red and obstacles are black. Parameter values are *p*_1_ = 1, *d*_1_ = 0.5, *m*_1_ = 5, *p*_2_ = *d*_2_ = *m*_2_ = 0, 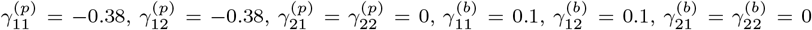, 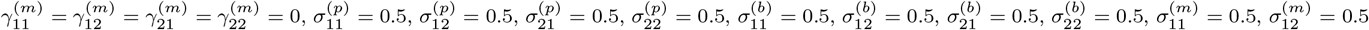, 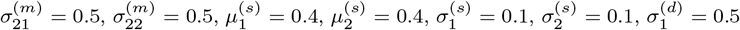, and 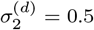

**Fig. 4.**
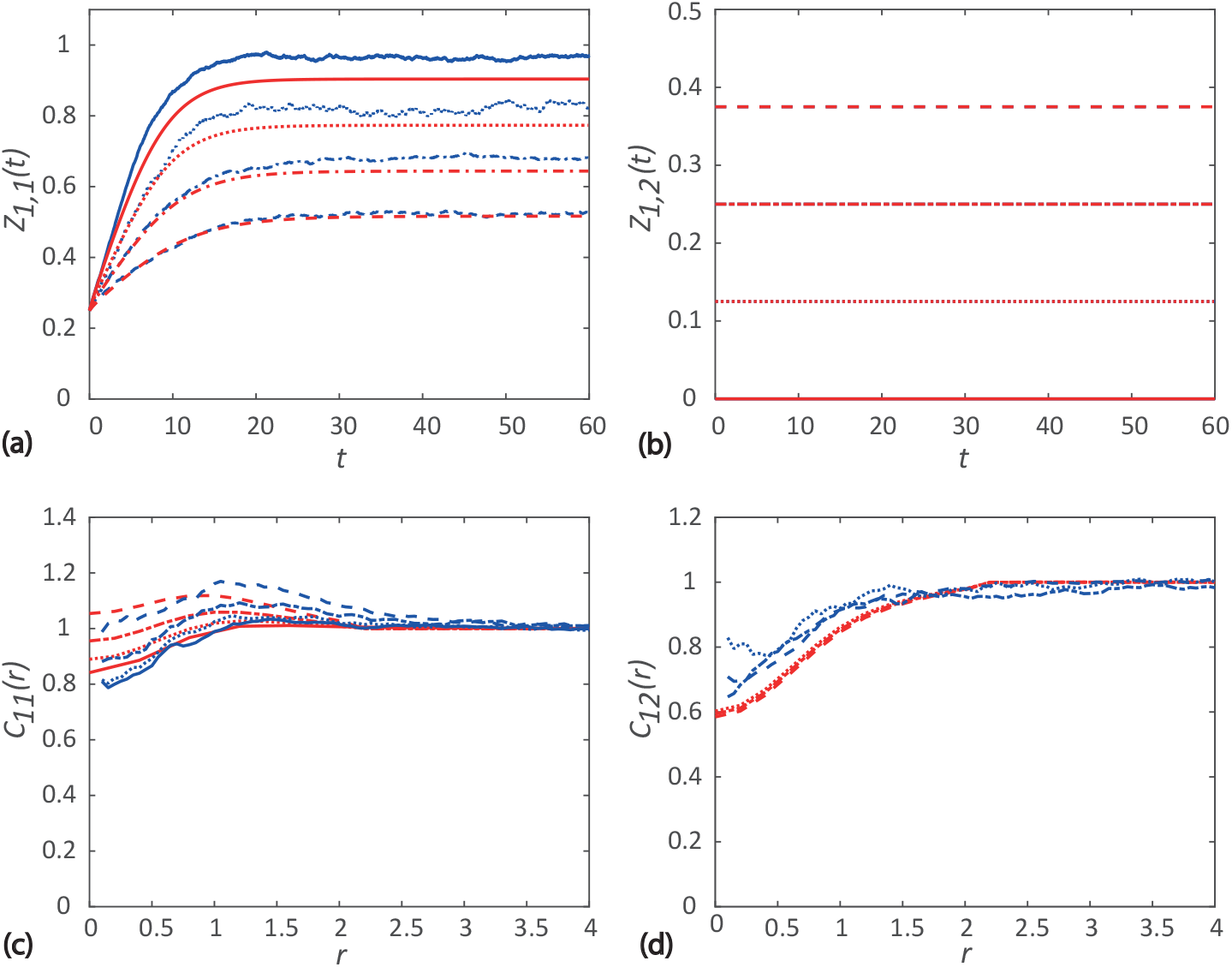
Comparison of spatial moment results and averaged data from 40 identically-prepared realizations of the IBM. Results in a show the evolution of the first spatial moment for the agents, *Z*_1,1_ (*t*). Various results are superimposed for different obstacle densities of obstacles: *Z*_1,2_(0) = 0/400 (solid), *Z*_1,2_(0) = 50/400 (dotted), *Z*_1,2_(0) = 100/400 (dash-dotted) and *Z*_1,2_(0) = 150/400 (dashed). Results in b show the constant first spatial moment for the obstacles. Results in c-d show the second spatial moment of agents expressed in terms of *C*_11_(*r*) and *C*_12_(*r*), respectively. Both PCFs are given at *t* = 60. The curves in red correspond to results from the spatial moment dynamics model, whereas curves in blue correspond to averaged data from the IBM. Parameter values are *p*_1_ = 1, *d*_1_ = 0.5, *m*_1_ = 5, *p*_2_ = *d*_2_ = *m*_2_ = 0, 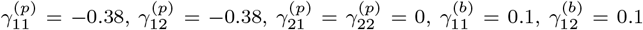, 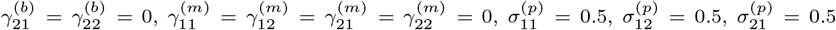, 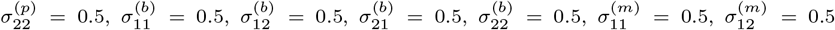, 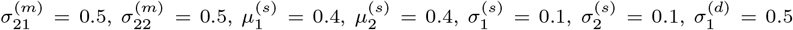, and 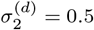.

Results in Figure 3(a)-(d) show the evolution of the population of agents and the associated spatial structure in the absence of obstacles. Overall we see that the initially small population of agents increases with time until there is a balance of net proliferation and death, leading to the formation of a steady density of agents at later time. For this choice of parameters we observe the formation of a regular spatial pattern. The main reason for the emergence of the regular pattern is the choice of a positive bias strength, 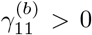 which means that agents tend to move away from regions of high density. The regular nature of the distribution of agents is evident in the auto-correlation PCF as we see that *C*_11_(*r*) < 1 over relatively short distances, *r*.

The incorporation of obstacles in the system leads to a reduction in the long time steady density of agents, as shown in Figure 4(a). In general we see that the more obstacles present, the smaller the steady state density of agents. This results makes intuitive sense as the presence of obstacles in the system increases the role of agent-obstacle interactions, which acts to reduce the net proliferation rate. These results also show that the accuracy of the spatial moment prediction increases with the density of obstacles. Results in Figure 4(b) show the time evolution of density of obstacles and we see the expected result that the density remains constant. However, the presence of obstacles influences the spatial structure of the population of agents by contributing to the directional bias. Since the obstacles are stationary, the obstacles prevent agents residing in certain regions of the domain. As we increase the obstacle density, we observe a progressive shift from the long time regular spatial pattern of agents over short distances when there are no obstacles present, to a more clustered long time pattern of agents as the obstacle density increases. The auto-correlation PCF for each of the cases shown in Figure 4(c) illustrates this transition. The cross-correlation PCF between agents and obstacles appears to be less sensitive to the density of obstacles than the auto-correlation PCF. For all cases we see that *C*_12_(*r*) < 1 over relatively short distances, r, for all the cases considered, indicating regular spatial pattern of agents and obstacles.

### 5.2 Effect of varying the obstacle size

Next, we explore how variations in obstacle size influence the spatial structure and dynamics of the agent subpopulation. The notion of obstacle size is incorporated into the model by varying the spatial extent of the interaction of obstacles, 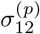 and 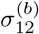. We assume that larger obstacles interact with agents over a greater distance than smaller obstacles. Therefore, we vary the size of the obstacles and examine how this impacts the evolution of the density of agents, and the spatial structure. We consider a total population composed of agents and obstacles with initial population sizes, *N*_1_ (0) = *N*_2_(0) = 100, respectively, as shown in Figure 5. We then consider increasing the obstacle size, shown from left-to-right in Figure 5, where we have 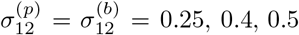, and 0.6, respectively. By repeating the stochastic simulations in Figure 5 many times, we can calculate the density of obstacles, the density of agents, the auto-correlation PCF and the cross-correlation PCF, as shown in Figure 6.

**Fig. 5.**
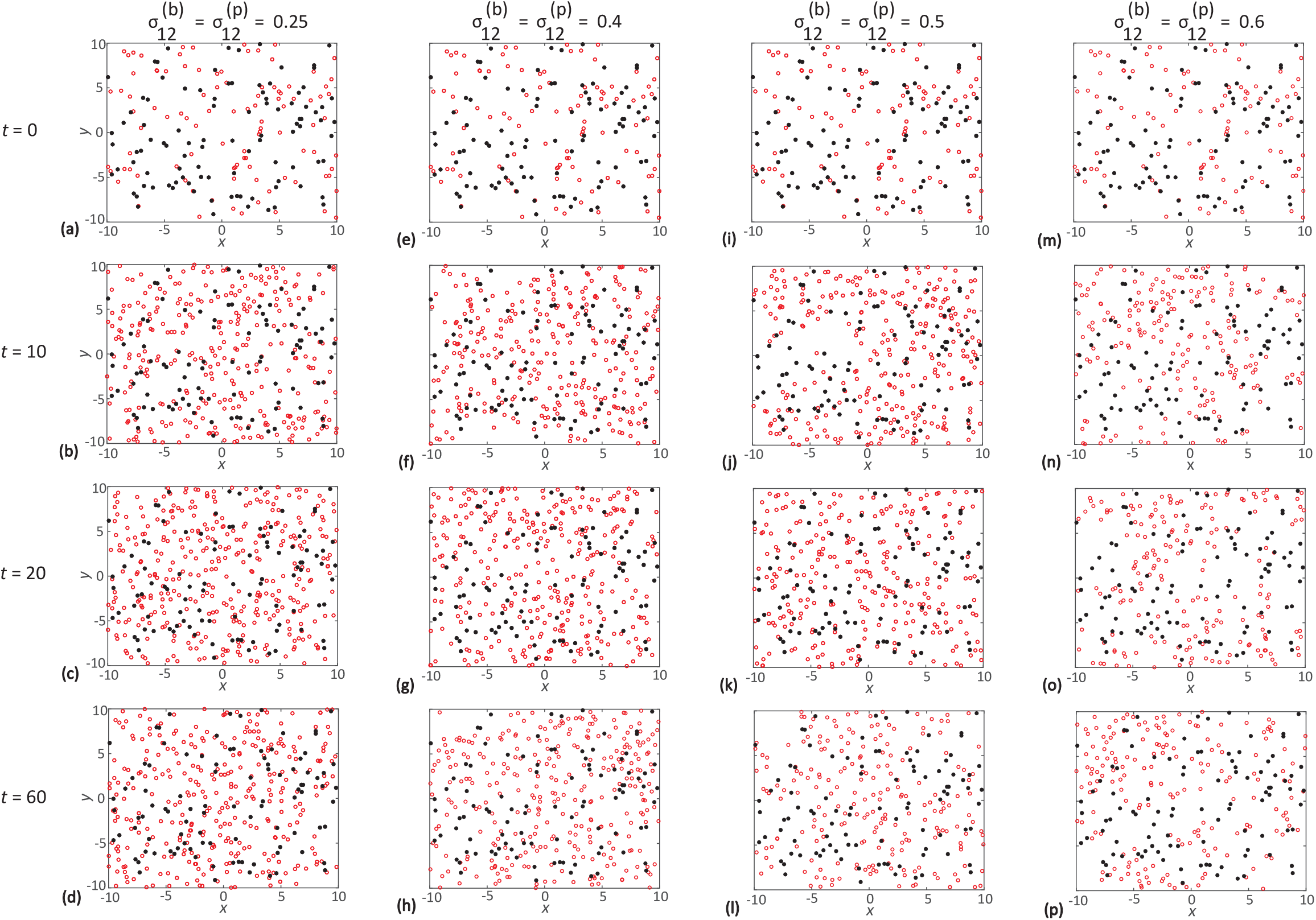
IBM snapshots showing the impact of increasing obstacle size. Each row shows a snapshot of the IBM at *t* = 0, 10, 20 and 60, respectively. Results in **a-d, e-h, i-l** and **m-p** show the evolution of a population of agents where 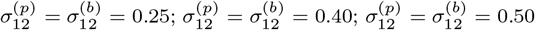; and 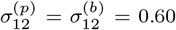, respectively. In each case the initial number of obstacles is 100. In all cases agents are red and obstacles are black. Parameter values are *p*_1_ = 1, *d*_1_ = 0.5, *m*_1_ = 5, *p*_2_ = *d*_1_ = *m*_2_ = 0, 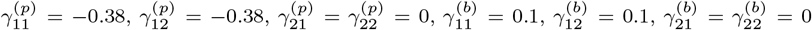, 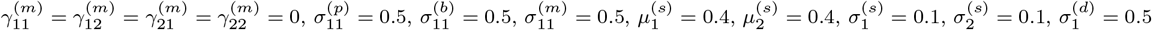, and 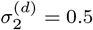.

**Fig. 6.**
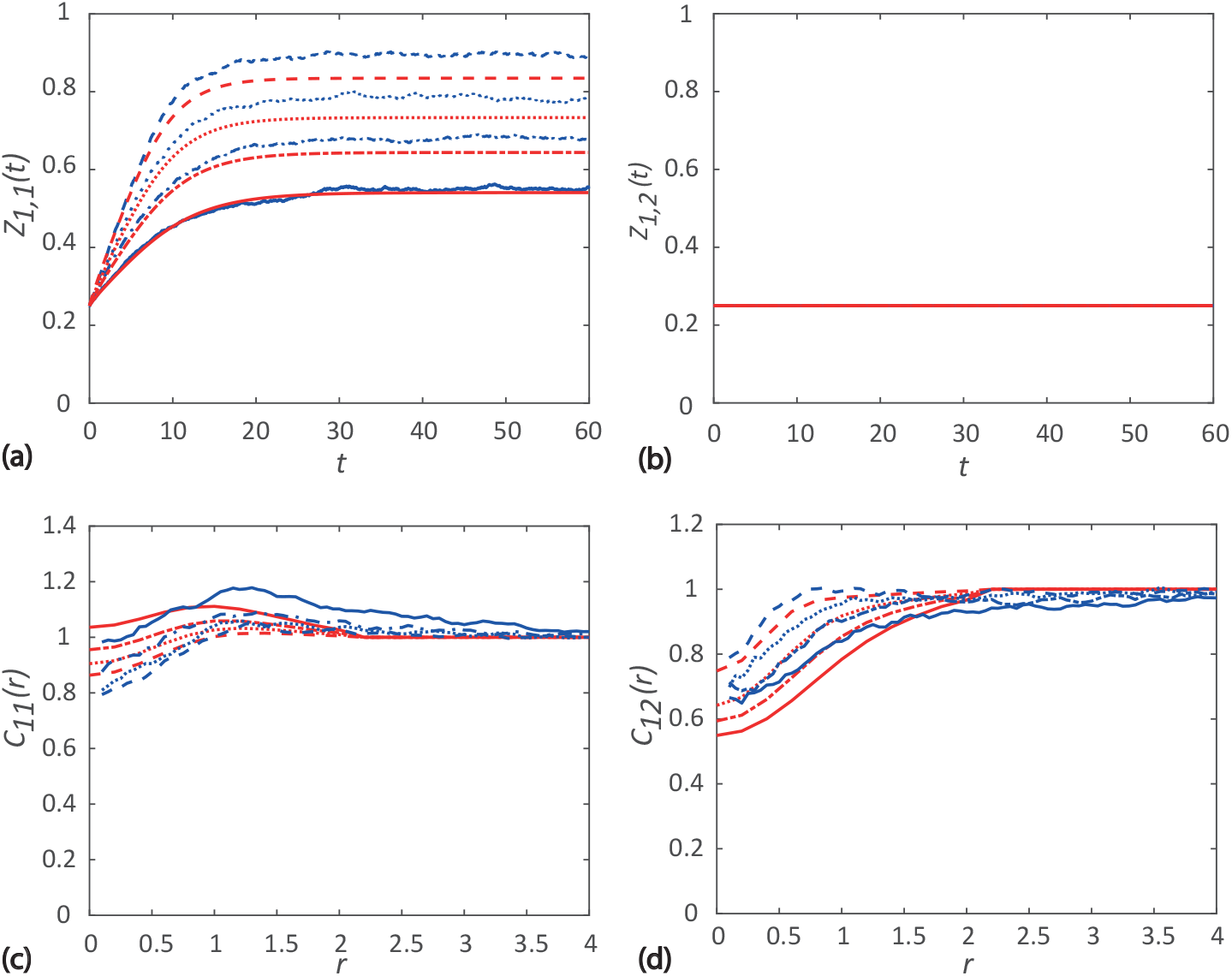
Comparison of spatial moment results and averaged data from 40 identically-prepared realizations of the IBM. Results in **a** show the evolution of the first spatial moment for agents, *Z*_1,1_ (*t*). Various results are superimposed for different obstacle size: 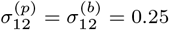 (dashed); 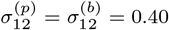 (dotted);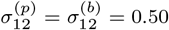 (dash-dotted); and 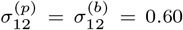 (solid). Results in **b** show the constant first spatial moment for the obstacles. Results in **c-d** show the second spatial moment of agents expressed in terms of *C*_11_(*r*) and *C*_12_(*r*), respectively. Both PCFs are given at *t* =60. **d** The curves in red correspond to results from the spatial moment dynamics model, whereas curves in blue correspond to averaged data from the IBM. Parameter values are *p*_1_ = 1, *d*_1_ = 0.5, *m*_1_ = 5, *p*_2_ = *d*_2_ = *m*_2_ = 0, 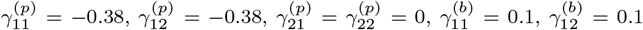, 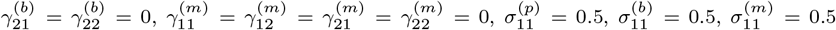, 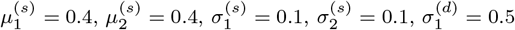, and 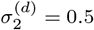.

Results in Figure 6(a)-(b) show the time evolution of the density of agents and the density of obstacles in each of the four cases considered. As the size of obstacles increase, we observe a decrease in the long time steady agent density. The wider range of interactions for the larger obstacles enables them to influence the proliferation rate of more distant agents. Even though the obstacle density is same, the proliferation rate of more agents reduces in the presence of large obstacles, leading to a reduced long time density of agents.

Figure 5 shows the snapshot of the spatial structure of population for each case considered and corresponding auto-correlation and cross-correlation PCFs are given in Figure 6(c)-(d). The population consisting of small obstacles shows a small-scale regular spatial pattern of agents. Due to the narrow interaction range of the smaller obstacles, a relatively small number of agents are affected by the obstacles. However, the agents are also subject to a repulsive bias from other agents, which results in a small scale regular spatial pattern of agents for this choice of parameters. The cross-correlation PCF is less than unity over short displacements, thereby suggesting regular spatial structure between agents and obstacles, but the effects are less pronounced than the spatial patterns between agents. As the obstacle size increases, the agent population became less regular, and the case we consider with the largest, 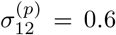, leads to agent clustering over short distances.

### 5.3 Effect of varying the obstacle interaction strength

Finally, we explore how variations in interaction strength, and the nature of the interactions between agents and obstacles, influence the spatial structure and the dynamics of the agent population. We examine the influence of both attractive interactions, as well as repulsive interactions. Similar to the previous simulations in Figure 5, we consider a random initial spatial distribution of agents and obstacles with initial population size *N*_1_(0) = *N*_2_(0) = 100. We then vary the strength of interactions, 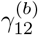, between 0.4 and −0.2. This means that we consider both attractive and repulsive interactions between agents and obstacles in this suite of simulations. Snapshots from the IBM are shown in Figure 7, and by repeating these stochastic simulations many times, we can calculate the density of obstacles, the density of agents, the auto-correlation PCF and the cross-correlation PCF, as shown in 8.

**Fig. 7.**
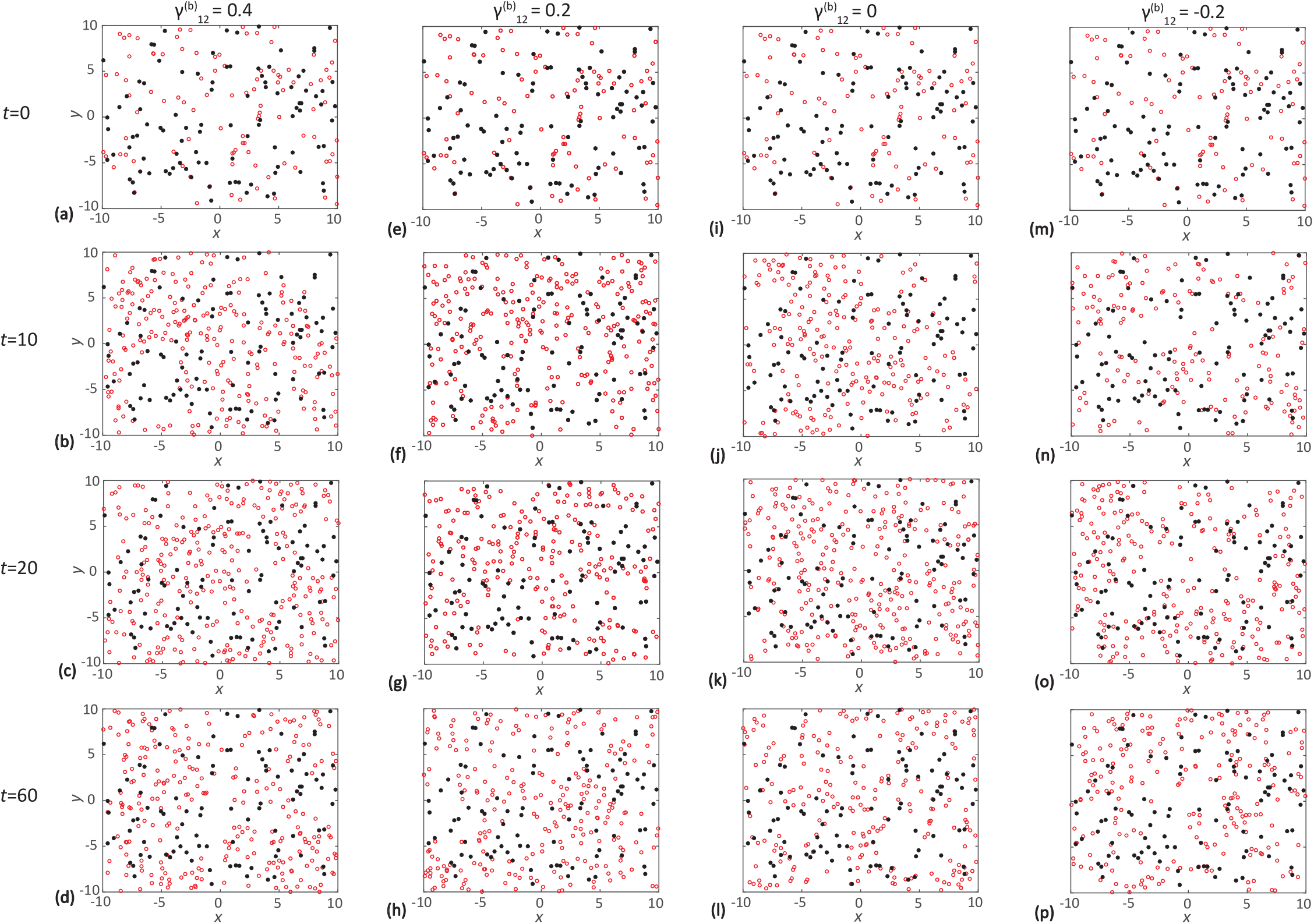
IBM snapshots showing the impact of increasing obstacle size. Each row shows a snapshot of the IBM at *t* = 0, 10, 20 and 60, respectively., **e-h, i-l** and **m-p** show the evolution of a population of agents where 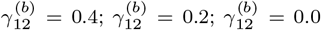; and 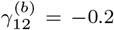, respectively. In each case the initial number of obstacles is 100. In all cases agents are red and obstacles are black. Parameter values are *p*_1_ = 1, *d*_1_ = 0.5, *m*_1_ = 5, *p*_2_ = *d*_2_ = *m*_2_ = 0, 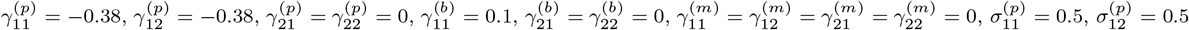, 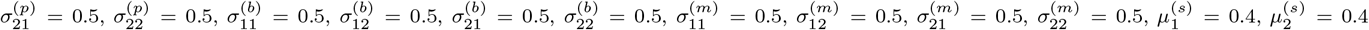, 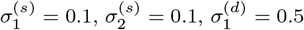, and 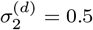.

Results in Figure 7(a)-(d) show the spatial structure arising when there is a relatively strong repulsive bias between the obstacles and agents. This repulsion means that agents tend to move away from the obstacles, leading to the formation of a regular spatial structure among agent-obstacle pairs. The PCFs, given in Figure 8(c)-(d), are consistent with this as we have *C*_12_(*r*) < 1 over small distances. Here we note that the repulsive interactions between agents are sufficiently strong to counteract the short-range dispersal of agents. Hence we also observe a regular spatial pattern among agents over short distances. Results in Figure 7(e)-(h) and Figure 7(i)-(l) show a similar spatial pattern, but the effects are less pronounced due to the reduced repulsion. Results in Figure 7(m)-(p) are quite different since we have attraction between the agents and obstacles, and the agents are biased to move towards the obstacles.

**Fig. 8.**
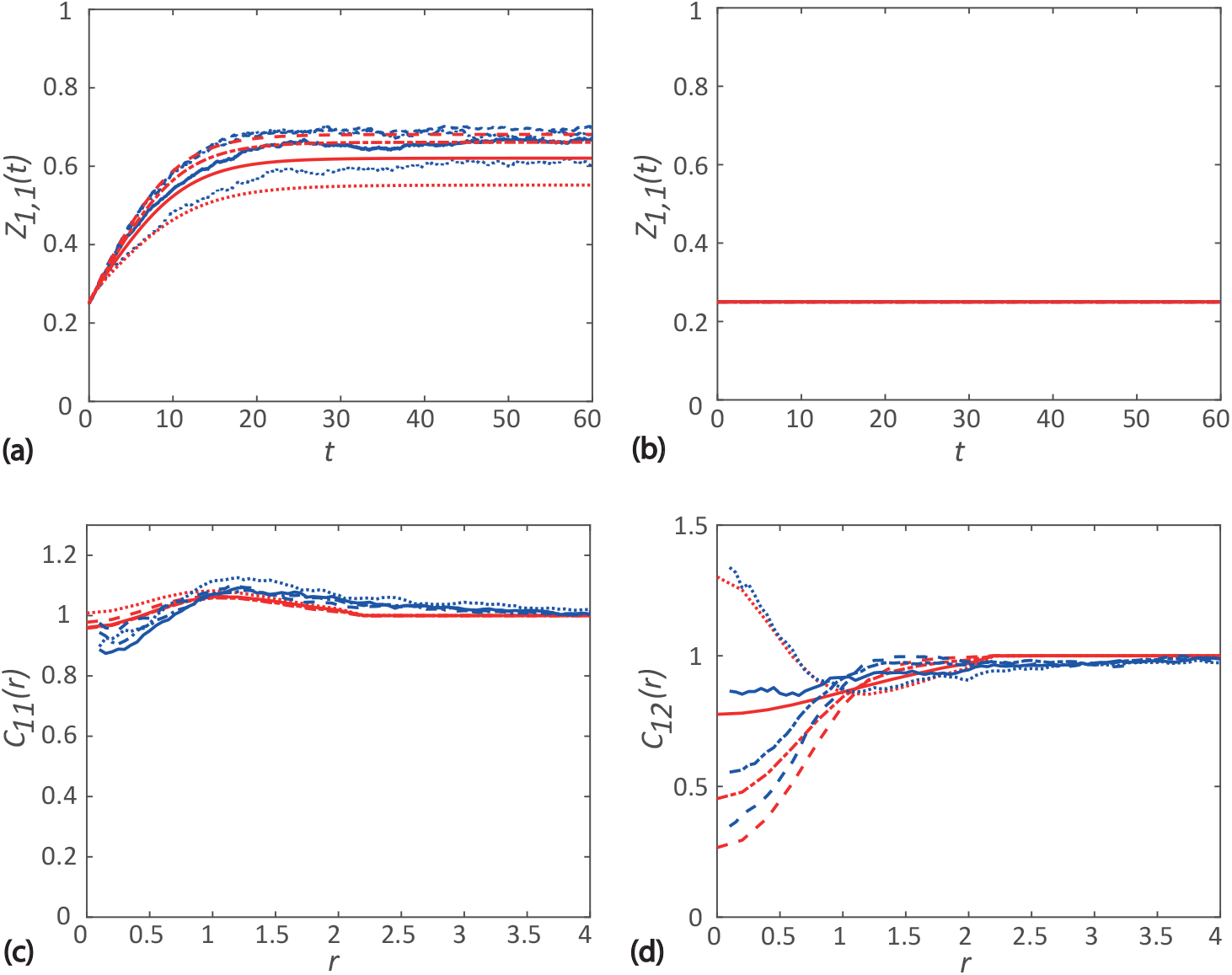
Comparison of spatial moment results and averaged data from 40 identically-prepared realizations of the IBM. Results in **a** show the evolution of the first spatial moment for agents, *Z*_1,1_(*t*). Various results are superimposed for different obstacle size: 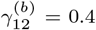 (dashed); 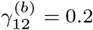 (dash-dotted); 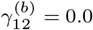 (solid); and 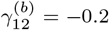 (dotted). Results in **b** show the constant first spatial moment for the obstacles. Results in **c-d** show the second spatial moment of agents expressed in terms of *C*_11(*r*_) and *C*_12_(*r*), respectively. Both PCFs are given at *t* = 60. **d** The curves in red correspond to results from the spatial moment dynamics model, whereas curves in blue correspond to averaged data from the IBM. Parameter values are *p*_1_ = 1, *d*1 = 0.5, *m*_1_ = 5, *p*_2_ = *d*_2_ = *m*_2_ = 0, 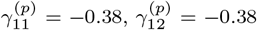, 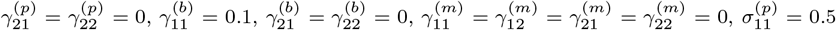, 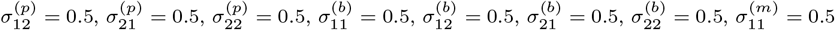, 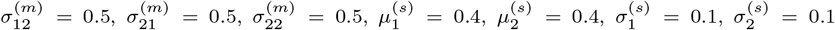, 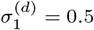, and 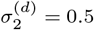.

Figure 8(a) shows the density dynamics of agents for each of the cases considered. The agent density decreases as the bias strength decreases and lowest when bias strength is negative. When the obstacle bias strength is negative corresponding to attractive obstacles, agents form clusters around obstacles. Since a large number of agents present at short distances, the proliferation rate of agents in clusters reduces significantly. Hence the population size increases more slowly than the case where obstacles are repulsive. Figure 8(b) shows the constant density of obstacles in each of the four cases.

## 6 Conclusion

In this work, we develop an IBM describing multi-species neighbour-dependent birth-death-movement processes, and we derive a continuum approximation of the stochastic dynamics using a spatial moment framework. We incorporate various processes such as neighbour-dependent directional bias from multiple subpopulations, and crowding effects such as contact inhibition of proliferation and contact inhibition of motility. We use this general framework to explore the case where one subpopulation is stationary and non-proliferative, and we treat this subpopulation as acting like biological obstacles in an *in vivo* environment. This framework allows us to explore how different properties of obstacles such as density, size and interaction strength influence the dynamics and emergence of spatial structure in the population.

Repeated simulations of the IBM are computationally inexpensive when the total number of individuals in the population is relatively small. However, such repeated simulations become computationally prohibitive as the population size increases. A useful feature of the spatial moment dynamics approximation is that the computational overhead is independent of the population size. Furthermore, if we were to explore the parameter space using the IBM to understand how variations in the interaction strength and the spatial extent of interactions lead to the emergence of spatial patterns, we would need to consider a large number of repeated stochastic simulations for each point in the parameter space. As we have shown in Figure 4 and Figure 6, simulations with certain parameter values lead to larger populations than other parameter combinations and the computational overhead associated with the IBM to describe these situations with larger population size could be significant. Under these situations, it might be more convenient to explore the parameter space using the spatial moment approximation since the computational cost is independent of the final population size.

Some previous models of cell movement treat individuals as hard spheres with a fixed radius to incorporate volume exclusion effects (e.g. Bruna and Chapman 2012; Dyson and Baker 2015; Plank and Simpson 2012). The main assumption behind these models is that the individual cells occupy space in the domain and any events would lead to other individuals overlapping the same space are prohibited. Our modelling framework does not impose absolute restrictions on the locations an individual can occupy. Instead, the probabilistic motility of individuals along with the neighbour-dependent bias kernel mean that events that lead to individuals being very close are assigned a low probability. Specifying neighbourhood interaction through an interaction kernel can be thought of as a proxy model for deformable, non-spherical individuals. This is an advantage of our model over volume exclusion models because cells are known to deform in the vicinity of other cells or obstacles (Hu et al. 2007; Le Clainche and Carlier 2008; Simpson et al. 2010).

Overall we see that the details of the dynamics of the population of agents and the spatial structure predicted using many identically prepared realisations of the IBM is reasonably well approximated by the solution of the spatial moment model. Our results reveal some interesting features that are not obvious without careful consideration. For example, results in Figure 3–4 show that as we increase the obstacle density, we observe that the steady state density of agents decreases, as we might expect. However, when we find that the accuracy of the spatial moments prediction increases with the density of obstacles, which is not obvious. Through our exploration we see the emergence of interesting spatial patterns as we vary the properties of obstacles. For example, results in Figure 3–4 shows that the long time steady arrangement of agents is regular at short distances when there is a sufficiently low density of obstacles present. In contrast, we see a clustered arrangement of agents at short distances when there is a sufficiently large density of obstacles present.

While our modelling framework is relatively general, there are many ways that it could be extended. For example, in this work we consider a spatially uniform initial distribution of individuals, and this kind of simulation is relevant to study certain cell biology experiments, such as a *proliferation assay.* However, other kinds of experiments, such as *scratch assays,* are initiated by considering an initial density of cells that varies spatially. To deal with this generalisation, both the IBM and our analysis needs modification. Furthermore, in this study we always consider stationary obstacles. However, in some applications it is thought that mobile obstacles are more relevant (Wedemeier et al. 2009), and this could be dealt with by setting the obstacle motility rate to be positive. Previous studies in ecology focusing on the birth-death processes of sessile organisms such as plants, consider fixed points in space corresponding to the locations of maximum resource availability in the habitats with an associated kernel (North and Ovaskainen 2007). Evaluating and summing the kernel over all these points leads to an environmental surface which measures the quality of habitats in the context of the evolution of dispersal. We note that the neighbour-dependent birth-death-movement model presented here can be extended to include the dispersal dependent upon densities of conspecifics or heterospecifics, or on habitat quality for both sessile and motile organisms (North et al. 2011). We leave these extensions for future consideration.

## Acknowledgements

This work is supported by the Australian Research Council (DP170100474). MJP is partly supported by Te Punaha Matatini, a New Zealand Centre of Research Excellence. We thank the anonymous referee for their helpful suggestions.

## References

1. Agnew DJG, Green JEF, Brown TM, Simpson MJ, Binder BJ (2014) Distinguishing between mechanisms of cell aggregation using pair-correlation functions. J Theor Biol 352: 16–23.

2. Bajenoff M, Egen JG, Koo LY, Laugier JP, Brau F, Glaichenhaus N, Germain RN (2006) Stromal cell networks regulate lymphocyte entry, migration, and territoriality in lymph nodes. Immunity 25(6): 989–1001.

3. Baker RE, Simpson MJ (2010) Correcting mean-field approximations for birth-death-movement processes. Phys Rev E 82: 041905.

4. Barraquand F, Murrell DJ (2013) Scaling up predator-prey dynamics using spatial moment equations. Methods Ecol Evol 4(3): 276–289.

5. Binny RN, Plank MJ, James A (2015) Spatial moment dynamics for collective cell movement incorporating a neighbour-dependent directional bias. J R Soc Interface 12(106): 20150228.

6. Binny RN, Haridas P, James A, Law R, Simpson MJ, Plank MJ (2016a) Spatial structure arising from neighbour-dependent bias in collective cell movement. PeerJ 4: e1689.

7. Binny RN, James A, Plank MJ (2016b) Collective cell behaviour with neighbour-dependent proliferation, death and directional bias. Bull Math Biol 78(11): 2277–2301.

8. Bolker B, Pacala SW (1997) Using moment equations to understand stochastically driven spatial pattern formation in ecological systems. Theor Popul Biol 52(3): 179197.

9. Browning AP, McCue SW, Binny RN, Plank MJ, Shah ET, Simpson MJ (2018) Inferring parameters for a lattice-free model of cell migration and proliferation using experimental data. J Theor Biol. 437: 251–260.

10. Condeelis J, Segail JE (2003) Intravital imaging of cell movement in tumours. Nat Rev Cancer 3(12): 921–930.

11. Dini S, Binder BJ, Green JEF (2018) Understanding interactions between populations: Individual based modelling and quantification using pair correlation functions. J Theor Biol 439: 50–64.

12. Dyson L, Baker RE (2015) The importance of volume exclusion in modelling cellular migration. J. Math. Biol. 71(3): 691–711.

13. Edelstein-Keshet L (2005) Mathematical Models in Biology (Classics in Applied Mathematics). Society for Industrial and Applied Mathematics, New York.

14. Ellery AJ, Simpson MJ, McCue SW, Baker RE (2014) Characterising transport through a crowded environment with different obstacle sizes. J Chem Phys 140: 054108.

15. Ellery AJ, Baker RE, McCue SW, Simpson MJ (2016) Modelling transport through an environment crowded by a mixture of obstacles of different shapes and sizes. Physica A 449: 74–84.

16. Finkelshtein D, Kondratiev Y, Kutoviy O (2009) Individual based model with competition in spatial ecology. SIAM J Math Anal 41(1): 297–317

17. Friedl P, Wolf K (2003) Tumour-cell invasion and migration: diversity and escape mechanisms. Nat Rev Cancer 3(5): 362–374.

18. Ghosh SK, Cherstvy AG, Grebenkov DS, Metzler R (2016) Anomalous, non-Gaussian tracer diffusion in crowded two-dimensional environments. New J Phys 18: 013027.

19. Gillespie DT (1977) Exact stochastic simulation of coupled chemical reactions. J Phys Chem 81: 2340–2361.

20. Hansen MM, Meijer LH, Spruijt E, Maas RJ, Rosquelles MV, Groen J, Heus HA, Huck WT (2016) Macromolecular crowding creates heterogeneous environments of gene expression in picolitre droplets. Nat Nanotechnol 11: 191–197.

21. Harley BA, Kim HD, Zaman MH, Yannas IV, Lauffenburger DA, Gibson LJ (2008) Microarchitecture of three-dimensional scaffolds influences cell migration behaviour via junction interactions. Biophys J 95(8): 4013–4024.

22. Hu K, Ji L, Applegate KT, Danuser G, Waterman-Storer CM (2007) Differential transmission of actin motion within focal adhesions. Science 315: 111–115.

23. Jin W, McCue SW, Simpson MJ (2018) Extended logistic growth models for heterogeneous populations. J Theor Biol 445: 51–61.

24. Johnston ST, Shah ET, Chopin LK, McElwain DLS, Simpson MJ (2015) Estimating cell diffusivity and cell proliferation rate by interpreting IncuCyte ZOOMTM assay data using the Fisher-Kolmogorov model. BMC Syst Biol 9: 38.

25. Keller EF, Segel LA (1971) Model for chemotaxis. J Theor Biol 30: 225–234.

26. Kurosaka S, Kashina A (2008) Cell biology of embryonic migration. Birth Defects Res Part C: Embryo Today 84(2): 102122.

27. Law R, Dieckmann U (2000) A dynamical system for neighbourhoods in plant communities. Ecology 81: 2137–2148.

28. Law R, Murrell DJ, Dieckmann U (2003) Population growth in space and time: Spatial logistic equations. Ecology 84: 252–262.

29. Le Clainche C, Carlier M (2008) Regulation of actin assembly associated with protrusion and adhesion in cell migration. Physiol Rev 88(2): 489–513.

30. Lewis MA (2000) Spread rate for a nonlinear stochastic invasion. J Math Biol 41: 430–454.

31. Martin P (1997) Wound healing-aiming for perfect skin regeneration. Science 276: 75–81.

32. Middleton AM, Fleck C, Grima R (2014). A continuum approximation to an off-lattice individual-cell based model of cell migration and adhesion. J Theor Biol. 359: 220–232.

33. Murray JD (1989) Mathematical biology. Springer, New York.

34. Murrell DJ, Dieckmann U, Law R (2004) On moment closures for population dynamics in continuous space. J Theor Biol 229: 421–432.

35. Murrell DJ (2005) Local spatial structure and predator-prey dynamics: counterintuitive effects of prey enrichment. Am Nat 166: 354–367.

36. North A, Ovaskainen O (2007) Interactions between dispersal, competition, and landscape heterogeneity. Oikos 116(7): 1106–1119.

37. North A, Cornell SJ, Ovaskainen O (2011) Evolutionary responses of dispersal distance to landscape structure and habitat loss. Evolution 65(6): 1739–1751.

38. Ovaskainen O, Cornell SJ (2006) Asymptotically exact analysis of stochastic metapopulation dynamics with explicit spatial structure. Theor Popul Biol 69(1): 13–33.

39. Ovaskainen O, Finkelshtein D, Kutoviy O, Cornell SJ, Bolker B, Kondratiev Y (2014) A general mathematical framework for the analysis of spatiotemporal point processes. Theor Ecol 7(1): 101–113.

40. Plank MJ, Law R (2015) Spatial point processes and moment dynamics in the life sciences: A parsimonious derivation and some extensions. Bull Math Biol 77: 586–613.

41. Plank MJ, Simpson MJ (2012) Models of collective cell behaviour with crowding effects: comparing lattice based and lattice-free approaches. J R Soc Interface 9: 2983–2996.

42. Raghib M, Hill NA, Dieckmann U (2011) A multiscale maximum entropy moment closure for locally regulated space-time point process models of population dynamics. J Math Biol 62: 605–653.

43. Simpson MJ, Towne C, McElwain DLS, Upton Z (2010) Migration of breast cancer cells: Understanding the roles of volume exclusion and cell-to-cell adhesion. Phys Rev E 82: 041901.

44. Simpson MJ, Binder BJ, Haridas P, Wood BK, Treloar KK, McElwain DLS, Baker RE (2013) Experimental and modelling investigation of monolayer development with clustering. Bull Math Biol 75: 871–889.

45. Simpson MJ, Plank MJ (2017) Simplified calculation of diffusivity for a lattice-based random walk with a single obstacle. Results Phys 7: 3346–3348.

46. Smith S, Cianci C, Grima R (2017) Macromolecular crowding directs the motion of small molecules inside cells. J R Soc Interface 14: 20170047.

47. Sun M, Zaman MH (2017) Modelling, signaling and cytoskeleton dynamics: integrated modelling-experimental frameworks in cell migration. WIREs Syst Biol Med 9: e1365.

48. Tan C, Saurabh S, Bruchez MP, Schwartz R, LeDuc P (2013) Molecular crowding shapes gene expression in synthetic cellular nanosystems. Nat Nanotechnol 8: 602–608.

49. Tobin P, Bjornstad ON (2003) Spatial dynamics and cross-correlation in a transient predator-prey system. J Animal Ecology 72: 460–467.

50. Wedemeier A, Merlitz H, Langowski J (2009) Anomalous diffusion in the presence of mobile obstacles. Europhys Lett 88: 38004.

51. Welch MD (2015) Cell migration, freshly squeezed. Cell 160: 581–582.

52. Zaman MH, Trapani LM, Sieminski AL, Mackellar D, Gong H, Kamm RD, Wells A, Lauffenburger DA, Matsudaira P (2006) Migration of tumor cells in 3D matrices is governed by matrix stiffness along with cell-matrix adhesion and proteolysis. Proc Natl Acad Sci USA 103: 10889–10894.

